# Genomic Repeat Element Analyzer for Mammals (GREAM)

**DOI:** 10.1101/003111

**Authors:** Darshan S Chandrashekar, Poulami Dey, Kshitish K Acharya

**Author notes:** To whom correspondence to be sent.

## Abstract

Background: Understanding the mechanism behind the transcriptional regulation of genes is still a challenge. Recent findings indicate that the genomic repeat elements (such as LINES, SINES and LTRs) could play an important role in the transcription control. Hence, it is important to further explore the role of genomic repeat elements in the gene expression regulation, and perhaps in other molecular processes. Although many computational tools exists for repeat element analysis, almost all of them simply identify and/or classifying the genomic repeat elements within query sequence(s); none of them facilitate identification of repeat elements that are likely to have a functional significance, particularly in the context of transcriptional regulation.

Result: We developed the ‘Genomic Repeat Element Analyzer for Mammals’ (GREAM) to allow gene-centric analysis of genomic repeat elements in 17 mammalian species, and validated it by comparing with some of the existing experimental data. The output provides a categorized list of the specific type of transposons, retro-transposons and other genome-wide repeat elements that are statistically over-represented across specific neighborhood regions of query genes. The position and frequency of these elements, within the specified regions, are displayed as well. The tool also offers queries for position-specific distribution of repeat elements within chromosomes. In addition, GREAM facilitates the analysis of repeat element distribution across the neighborhood of orthologous genes.

Conclusion: GREAM allows researchers to short-list the potentially important repeat elements, from the genomic neighborhood of genes, for further experimental analysis. GREAM is free and available for all at http://resource.ibab.ac.in/GREAM/

## Introduction

Differential regulation of genes is crucial for the development of specific cell types, tissues and the onset/maintenance of specific physiological conditions in multicellular eukaryotes. Efforts are on, by many scientists, to decode the mechanisms of such regulation.

From the time Barbara McClintock reported about transposable elements [Fedoroff, 2001], many researchers may have wondered about a possible influence of such elements on gene expression in broader contexts. Repeat elements constitute about 69% of the human genome [de Koning, 2011]. However, until recent years, their role in the regulation of gene expression in specific cell type/physiological conditions was not successfully explored.

Transposition within a protein coding region can either harm the cellular functions or benefit by generating the transcript diversity via novel splice sites or regulation signals [Cowley, 2013]. Transcriptional interference of retro-elements has been indicated to be one of the contributing factors for shaping the human genome in terms of distribution of the transposable elements and protein-coding genes [Mourier and Willerslev, 2008]. Possible involvement of repeat elements in the transcription regulation, via defined chromatin loops, was suggested earlier based on the transposon-enrichment in scaffold/matrix associated regions [Jordan, 2003]. In fact, the genomic repeat elements are now indicated to have a significant role in the gene regulatory networks [Estécio, 2010; Lynch, 2011; Kunarso, 2010].

Hence, there is a need for larger-scale exploration of the role of repeat elements in the transcription regulation, particularly for the clusters of co-expressed genes. In this context, a preliminary *in silico* screening of all genomic repeat elements is particularly important to short list the potentially important ones. But a suitable bioinformatics-software is not available for such screening and/or analysis. Even though there are many tools already developed for repeat element detection [Lerat, 2010], none of them allow identification of over-represented repeat elements in the neighborhood region of a given set of genes. Among available resources, only TranspoGene [Levy, 2008] allows query with multiple genes and reveals repeat elements from the intronic, exonic and proximal promoter region of the gene. However, TranspoGene also has limitations (see table 1). For example, it lists repeat elements located only in a short (up to 250bp) upstream region. It also does not summarize repeat element distribution across genes. Thus, it is currently not possible to identify over- or under-represented elements, particularly across larger neighbourhoods. Observation of such limitations prompted us to develop a new web server that enables users to analyze repeat element distribution in the neighborhood of co-expressed/co-regulated genes: ‘Genomic Repeat Element Analyzer for- Mammals’ (GREAM, http://resource.ibab.ac.in/GREAM).

**Table 1.** Comparison of GREAM with TranspoGene based on information content and output features.

| **Feature** | **GREAM** | **TranspoGene** |
| --- | --- | --- |
| Allows users to submit set of genes as input | Yes | Yes |
| Allows users to analyze repeat element distribution in the neighborhood of gene start site. | Yes (up to 3MB upstream and 3MB downstream from gene start site) | Yes (up to 250 bp upstream and the entire length of gene) |
| Identifies statistically over-represented repeat elements in the neighborhood of given set of genes | Yes | No |
| Number of mammalian species for which analysis can be performed | 16 | 2 |
| Allows user to analyze repeat element distribution in the orthologous genes | Yes | No |
| Categorize repeat elements based on their location within gene(exon/intron region) | Yes | Yes |
| Repeat element information last updated | September 2013 | 2008 |

## Results

GREAM is developed using PERL-CGI. The workflow of GREAM is schematically represented in figure 1.

**Figure 1.**
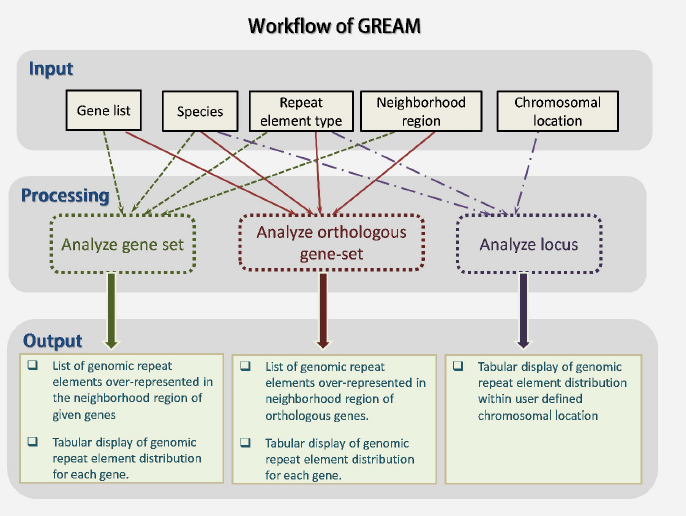
Workflow of the GREAM tool. The input parameters considered for different type of analysis and output obtained from the tool is mentioned

GREAM allows user to analyze repeat element distribution in the neighborhood of, (a) a set of genes from a species, through ‘Analyze gene-set’ feature; (b) a set of genes in multiple mammalian species, through ‘Analyze orthologous gene-set’ feature; or (c) a specific chromosomal location, through ‘Analyze locus’ feature.

### Input

While using ‘analyze gene set’ and to ‘analyze orthologous gene-set’ sections, users need to submit a gene list (either by pasting in text area provided or by uploading a text file) and an appropriate species. Gene list needs to be homogenous in terms of the identifiers; the genes in the list can be represented by the official gene symbols, NCBI identifier, Ensembl gene identifier, RefSeq mRNA identifier or the Unigene identifier. The length of the neighborhood regions can be identified in two ways: a) using the start site of the query gene as the central point or b) using the gene itself as center, i.e., choosing the region before the gene and after it. There is an option to limit the analysis to specific repeat element class as well. The ‘analyze locus’ section requires user to specify chromosomal location for chosen species.

There is an option to choose between the output-display in the web-interface or obtaining the output via email, as in some cases there may be a significant waiting required. For instance, if a user submits more than 100 genes to ‘analyze gene-set’ option, or more than 25 genes considering four or more species to ‘analyze orthologous gene-set’ option, GREAM will take a while to process, and an email-mode is recommended in such cases.

### Processing

#### Analyze gene-set

- The tool scans the gene list and converts all valid identifiers to official gene symbols using Ensembl 70 gene identifier track from BioMart [http://asia.ensembl.org/biomart/martview]. It then obtains gene start sites from NCBI Gene annotation track [ftp://ftp.ncbi.nih.gov/gene/].
- All repeat elements located in the user-defined regions are identified using repeat annotation track from NCBI [ftp://ftp.ncbi.nih.gov/gene/].
- The statistical significance of abundance of each repeat element in the specific regions around the query-genes, as compared to the expected frequency of occurrence across the genome, is assessed by using binomial probability.

#### Analyze orthologous gene-set

- Here, in addition to scanning and identifying valid genes, the tool uses in-house MySQL database to obtain orthologous genes (developed using NCBI HomoloGene Build 67 for seven mammalian species [*Homo sapiens*, *Mus musculus*, *Rattus norvegicus*, *Bos taurus*, *Canis familiaris*, *Pan troglodytes* and *Macaca mullata*].
- Distribution of repeat elements in the neighborhood region is analyzed not only for submitted gene lists, but also on their orthologous genes.
- The tool will identify repetitive elements that are over-represented in the neighborhood of the query genes, for the main species as well as for orthologous genes from selected species.

#### Analyze Locus

Based on species and chromosomal location provided, suitable repeat annotation track is used to analyze repeat element distribution in the locus mentioned. This search feature is currently functional for 15 species. The feature could not be enabled for *Loxodonta africana* and *Nomascus leucogenys* species for which the contig-assembly is incomplete.

### Output

Distribution of different classes of repeat elements within the neighborhood region of each gene submitted is shown in a tabular format; relative frequency of each element-class is also depicted as a histogram (generated via open flash chart [teethgrinder.co.uk/open-flash-chart/]).

The results for ‘Analyze gene-set’ and ‘Analyze orthologous gene-set’ includes an additional table that summarizes distribution of specific repeat elements across neighborhood region of submitted genes, along with its statistical significance (assessed through binomial probability, see methods) of occurrence in selected regions.

## Discussion

### Validation using a case study

To ensure the functionality of the newly developed tool we validated it using experimental results by Lynch et al [Lynch, 2011]. Their results revealed over-representation of transposable element MER20 in the neighborhood region of endometrial expressed genes; 200kb from before start and equal region after the end of genes was considered. We downloaded ‘processed human RNA-Seq data’ from GEO (GSE30708) and identified 1149 genes (having read count of >20 as mentioned in the study) up- or down-regulated in differentiated human endometrial tissue compared to undifferentiated human endometrial tissue. When analysis with GREAM was repeated considering 200kb upstream of gene start site and 200kb upstream of gene end site, MER20 was found to be significantly (P-value: 0.015) over-represented in neighborhood of 943 (82%) of these genes. GREAM also provided additional information. For example, of the total of 1332 repeat elements were found detected in the neighborhood of the query genes, 530 other repeat elements were also found to be over represented at a statistical level of <=0.015.

### Utility of the tool

The above-mentioned case study illustrates the use of GREAM for short-listing genomic repeat elements around genes that may have a functional significance. Thus the tool can be used to short list repeat elements for experimental analysis. The following questions represent examples of other research problems that can be addressed using GREAM:

a. Which repetitive elements might be strongly associated with epigenetic regulation of gene promoters?
b. Are there similar/same repetitive elements enriched in the neighborhood of genes that show constitutive expression across multiple tissues and multiple organisms?
c. Is there an abundance of any specific type of repeat element in the neighborhood of chromosomal loci found to be frequently associated with a specific type of structural abnormality?
d. Are there repeat elements associated with a set of differentially transcribed genes? If yes, which ones, and are they conserved in cases of othologous genes?
e. Are there y-chromosome-associated repeat elements preserved across mammalian species and, if yes, in which regions?
f. Do genes near telomeres and/or centromeres have different type of repeat elements?
g. Are there any enriched repeat elements near the centromeres and/or telomeres, other than the standard repeats already known to be associated with these chromosomal components?

### Conclusions

The reported online tool, GREAM, would be useful for selecting important genomic repeat elements around specific genes or other chromosomal regions of interest. GREAM offers features that do not exist in any other already existing tool.

## Methods

### Information collection

Different gene identifiers such official gene symbol, Entrez Gene id, Ensembl Gene id, RefSeq mRNA id and UniGene id corresponding to all 16 mammalian species considered were downloaded from Ensembl BioMart (http://asia.ensembl.org/biomart/martview). Orthologous gene information was downloaded from NCBI HomoloGene Build 67 (ftp://ftp.ncbi.nih.gov/pub/HomoloGene/). Repeat annotation tracks of mammalian genome sequences obtained using RepeatMasker [http://www.repeatmasker.org/] were downloaded from NCBI Gene FTP (ftp://ftp.ncbi.nih.gov/gene/). Similarly gene-related information for every mammalian species considered was obtained from NCBI Gene FTP (ftp://ftp.ncbi.nih.gov/gene/).

### Estimating over-representation of repeat element

The over-representation of every repeat element is assessed by binomial probability [Ross, 2010]. We determined the frequency of occurrence of each repeat element within neighborhood regions (10KB, 20KB, 50KB, 100KB, 200KB, 500KB, 1MB, 2MB and 3MB) of all protein coding genes (as per NCBI gene database) for every mammalian species. When a user submits a set of genes (n) and observes occurrence of any repeat element within specified neighborhood regions of some of or all of these genes [k<=n], the significance of its occurrence is estimated using the following formula:

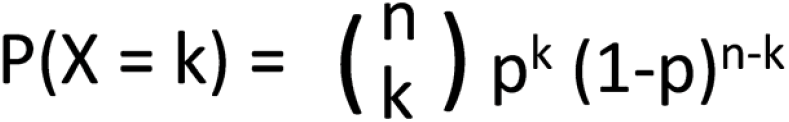

Where, P(X = k) is probability of occurrence of specific repeat element in the neighborhood of ‘k’ genes out of ‘n’ genes submitted, ‘p’ is probability of occurrence of specific repeat element obtained by population statistics (total number of protein coding genes having specific repeat element within neighborhood region / total number of protein coding genes in the genome)

If the calculated probability of observing repeat element in neighborhood regions of k genes is less than 0.05, it can be considered significant. The tool displays this probability value along with the details of ‘k’ genes.

### Data access

GREAM is freely available, both as online version and offline version. Users can download all PERL scripts, MySQL database, gene annotation and repeat annotation data from the GREAM site (http://resource.ibab.ac.in/GREAM/).

## Acknowledgements

The authors also thank Mr. Anish Kumar M, Mrs. Madhura Adavkar, Mr. Venkata Shiva Prasad and Ms. Pallavi Gupta, students from IBAB, for their efforts in data collection from different repositories and validation. This work was supported by an institutional grant to IBAB from Department of Electronics and Information Technology (DeitY), Government of India, under ‘Centre of Excellence for Research and Teaching in Bioinformatics'.

## Authors’ contributions

DSC wrote the PERL scripts, created the tool and performed the case study for validation. PD identified the statistical methods for analysis. KKA conceptualized the project and guided the progress.

